# Compositional analysis of yellow fever virus induced stress granules reveals a functional connection to mitochondrial homeostasis

**DOI:** 10.1101/2025.11.13.688208

**Authors:** Celia Jakob, Felix Streicher, Marta Lopez-Nieto, Lucy Eke, Diane Lee, Jinjun Wu, J. Paul Taylor, Ana M Matia-Gonzalez, Nolwenn Jouvenet, Nicolas Locker

## Abstract

The phase separation of biomolecules into so-called stress granules (SGs) allows the cell to tightly regulate translation activity in response to different stimuli, such as oxidative stress, starvation, or the recognition of non-cellular RNA. Recent reports suggest SGs induced during viral infection, may act as a crossroad between the cellular stress response and the activation of the innate immune response.

Here, we aimed to dissect the role of SGs in the context of yellow fever virus (YFV) infection. We found that YFV infection resulted in translational shut-off from 24 hpi on followed by the formation of SGs at 48 hpi, a delay potentially associated with the ability of the YFV capsid to inhibit SG formation, through an interaction with the major SG scaffolding protein G3BP1. To elucidate the role of YFV-induced SGs during infection, we inhibited SG assembly using a small-molecule inhibitor and find that SG formation does not influence viral replication. Uncovering the first proteome of virus-induced SGs, our compositional analysis revealed a specific enrichment of proteins associated with mitochondrial processes in YFV-induced SGs. Indeed, we show that YFV infection results in mitochondrial damage and dysfunction. Together, we propose that YFV-induced SGs may be involved not only in the modulation of cellular homeostasis but also in influencing mitochondrial functions.

**Author Summary:** Viruses impose a major burden on the infected host – from structural rearrangements needed to assemble replication complexes, to exploiting cellular energy resources and genetic rewiring associated with antiviral responses. The assembly of membrane-less organelles such as stress granules (SGs) enable cells to rapidly tune cellular processes upon sensing of stresses such as viruses. Moreover, the cell’s innate immune response is proposed to be regulated by SGs and in turn many viruses disrupt or highjack their components. Yet, the molecular basis for SG functions during infection remain ambiguous.

Here we investigated the interplay between yellow fever virus (YFV) infection and SGs. We demonstrate that infection with attenuated or pathogenic viruses result in the formation of SGs. Their compositional analysis reveal that they sequester mitochondrial proteins, correlating with altered mitochondrial functions during infection. This highlights a novel complex interplay between membrane-bound and membrane-less organelles which could present novel opportunities for antiviral therapies.

## Introduction

Stress granules (SGs) are biomolecular condensates that assemble by liquid-liquid phase separation and are involved in translational regulation and maintenance of cellular homeostasis (1). The formation of SGs has originally been described as part of the cellular intrinsic stress response (ISR), which can be activated by sensing oxidative stress, nutrient starvation or endoplasmic reticulum (ER) stress, as well as viral infections (1,2). Activation of the ISR typically results in the phosphorylation of eukaryotic initiation factor 2α (eIF2α) by kinases including HRI, PERK, GCN2 and PKR, thereby impeding the initiation of 5’ cap-dependent mRNA translation (1). This process is accompanied by the disassembly of the translation initiation complex. Subsequently, translationally silent messenger ribonucleoproteins (mRNPs) are captured by multifunctional SG proteins (e.g. G3BP1, TIA1 and Caprin1) and condensed into cytoplasmic SGs (3,4). SGs are typically structured into a solid core surrounded by a dynamic shell, enabling exchange of components with the cytoplasmic environment (5). When cellular stress is alleviated, the disassembly of SGs is regulated by post-translational modifications and the proteasomal degradation of SG proteins, freeing mRNAs for translation (1).

Viral infections lead to significant disruption of cellular homeostasis. For example, sensing of viral products, such as replication intermediates or proteins, via cellular pattern recognition receptors (PRRs) results in the activation of the innate immune response (IIR), accompanied by the upregulation of antiviral proteins, cytokines, and interferons (6). Furthermore, ongoing viral replication activates the ISR by sensing dsRNA, for example, via PKR, as well as ER and oxidative stress, which leads to translational shut- off (2). Interestingly, it has been proposed that SGs can concentrate diverse PRRs including RIG-I, MDA-5, or PKR (7–12). They have therefore been proposed to be at a crossroad integrating the ISR and the IIR pathways. However, the exact role of SGs in modulating the IIR or viral replication remains unclear. While some studies have shown that genetic disruption of SG formation results in reduced activation of the IIR and increased viral replication (7,8,11), other studies have reported that SG disruption results in an excessive immune response (12). The latter has been shown to be coupled to an increase in cell death. Accordingly, SGs can sequester executioner caspases, to promote cell survival (13). Despite these observations, recent studies have questioned the direct involvement of SGs in regulating the IIR, as they have failed to observe PRR co- localisation with SGs or altering SG formation did not influence innate immune activation (10,14–17). Nevertheless, while the role of SGs in the IIR remains controversial, studies focusing on SG formation during viral infection suggest that their resident proteins have a functional role beyond translational control (18,19). Consequently, many viruses have evolved strategies to prevent SG formation by inhibiting eIF2α phosphorylation or by cleaving or sequestering SG proteins (2). Furthermore, recent observations point to the existence of SG-like structures that exhibit anti-viral functions independently of translational shut-off (20). For example, during SARS-CoV-2, flavivirus and influenza virus infection, viral genomic or mRNA can be sequestered into condensates in a G3BP1- dependent manner (8,15). Similarly, viral infection can trigger the formation of SG-like condensates, which contain SG proteins but form independently of G3BP1. RNaseL- induced bodies (RLBs), for example, form upon sensing of viral RNA and are associated with the anti-viral RNA decay machinery (20,21). Although RLBs enrich for G3BP1, they lack typical SG proteins such as TIA-1 and form independently of eIF2α phosphorylation. Similarly, the formation of paracrine granules has been observed in uninfected bystander cells (20,22). These paracrine granules accumulate but do not require G3BP1/2, lacks eIFs, and protect bystander cells from infection. These studies highlight our limited understanding of the functional and compositional diversity of SGs and SG-like structures during viral infections.

Yellow Fever Virus (YFV) belongs to the *Orthoflaviviridae* family, which comprises single-stranded, positive-sense RNA viruses that typically possess a type I cap at the 5’ end of the viral genome (23). The viral genome therefore acts as mRNA and can be directly translated upon cell entry. Many orthoflaviviruses are transmitted to vertebrate hosts by arthropod vectors and can cause severe diseases ranging from flu-like symptoms to organ failure, as seen with YFV infection (24). Other prominent human pathogens in the *Orthoflavivirus* genus include Zika virus (ZIKV), West Nile virus (WNV), Japanese encephalitis virus (JEV) and dengue virus (DENV) (23). While it has been demonstrated that many of these can be sensed via PKR (25–29), conflicting results exist regarding the phosphorylation of eIF2α and subsequent translational shut-off during viral infection. For WNV, it has been shown that activation of the PKR-eIF2α axis is strain-specific and relates to early vRNA synthesis (28). For DENV and ZIKV, however, conflicting reports show diverse effects on the phosphorylation of eIF2α and translational shut-off (25–27,30). These differences seen in DENV and ZIKV studies may be linked to the different viral strains, cell lines and infection conditions used. Interestingly, infection with ZIKV, JEV, DENV and WNV all appear to inhibit SG formation (19,25,27,28,30–32). This has been proposed to be due to the ability of specific viral proteins to hijack SG proteins for their own replication, thereby sequestering them away from SG formation (27,31,32). Notably, despite yellow fever being one of the earliest viral diseases to be studied, the cellular stress response to YFV infection remains largely unexplored. A recent study showed that YFV is sensed by PKR and RIG-I (17). In contrast to other orthoflaviviruses, the authors observed the robust induction of SGs, which appeared to accumulate RIG-I. However, these SGs did not appear to modulate the innate immune response, as knocking down several SG proteins did not affect cytokine or interferon production upon infection. This leaves the functional relevance of these granules during YFV infection unknown.

Here, we aimed to elucidate the interplay between YFV infection and SG formation. To this end, we studied two different YFV strains: YFV/17D, a live-attenuated strain used for vaccination, and YFV/Asibi, a highly pathogenic wild-type strain (33). We found that both strains induced translational shut-off and robust SG formation at late stages of infection. These granules displayed features of canonical SGs, as they are sensitive to agents that stabilize translational complexes. We found that the delay in SG formation relative to the onset of translational shut-off, might relate to the ability of the viral capsid protein to inhibit SG assembly. In this regard we show, that the YFV capsid protein contains a binding motif for the major SG protein G3BP1, which is required for the capsid-G3BP1 interaction *in vitro*. To dissect the role of YFV-induced SGs during infection, we treated infected cells with a small-molecule inhibitor called G3Ib, that has been shown to inhibit SG assembly by binding to G3BP1. We found that inhibition of SGs does not influence YFV replication *per se*. Performing compositional analysis, we revealed that virus-induced SGs are linked to mitochondrial function. Indeed, we observed that YFV infection induced mitochondrial damage associated with depolarization and increased ROS production. Overall, this suggests that YFV has evolved to tolerate SGs during infection to modulate mitochondrial functions.

## Results

### YFV infection induces translational shut-off and the formation of canonical stress granules

Previous studies have reported that infection with orthoflaviviruses, such as ZIKV, JEV and DENV, results in translational shut-off, while the formation of SGs appears to be suppressed (25–27,29–32). However, the effect of YFV on translational regulation has not yet been investigated. Therefore, we applied the ribopuromycylation (RPM) assay in Huh7 cells to measure *de novo* protein synthesis during infection with two different YFV strains: the attenuated vaccine strain YFV/17D and the highly pathogenic strain YFV/Asibi (33). To control for translational shut-off, we treated the cells with sodium arsenite (NaAs) for 30 minutes (mins). NaAs is known to induce oxidative stress, resulting in reduced translational activity (34). To control specifically for infection-induced translational shut- off events, we infected Huh7 cells with ZIKV for 24 hours (h) (25,27). Both, ZIKV infection and NaAs treatment resulted in strong translational shut-off (S1A-S1C Fig). At 24 h post-infection (hpi), cells infected with YFV/17D exhibited reduced translational activity, albeit not to the same extent as ZIKV-infected cells. Similarly, YFV/Asibi infection induced translational shut-off compared to mock-infected cells. Translational shut-off is typically accompanied by the formation of stress granules (SGs) (1). To assess whether YFV infection triggers SG formation, we performed staining against eIF3η, which is known to localise to SGs (3,4). While we observed robust SG formation in 98 % of the NaAs-treated cells, we observed only in 1 % of YFV/17D- and in 15 % of YFV/Asibi-infected cells the formation of SGs at 24 hpi (S1A Fig, Fig 1 B). Contrary to previous reports, we also observed SG formation in 65 % of ZIKV-infected cells. We speculated that different virus kinetics might result in delayed SG induction in YFV-infected cells. Therefore, we examined cells infected with YFV/17D and YFV/Asibi at 48 hpi for translational shut-off and SG formation. As observed at 24 hpi, translational activity in YFV-infected cells remained reduced at 48 hpi (Fig 1 A, C, D and S2A-S2B Fig). However, at this later time point, we also observed SG formation in 47 % of YFV/17D-infected cells and 59 % of YFV/Asibi-infected cells (Fig 1 B). A typical trigger for translational shut-off and SG assembly is the phosphorylation of eIF2α following the activation of eIF2α kinases, including PKR, which impairs the recycling of ternary complexes (1). In agreement with the observed translational shut-off and SG formation, infection with ZIKV, YFV/17D and YFV/Asibi resulted in robust eIF2α phosphorylation (S2C-S2D Fig).

**Fig 1.**
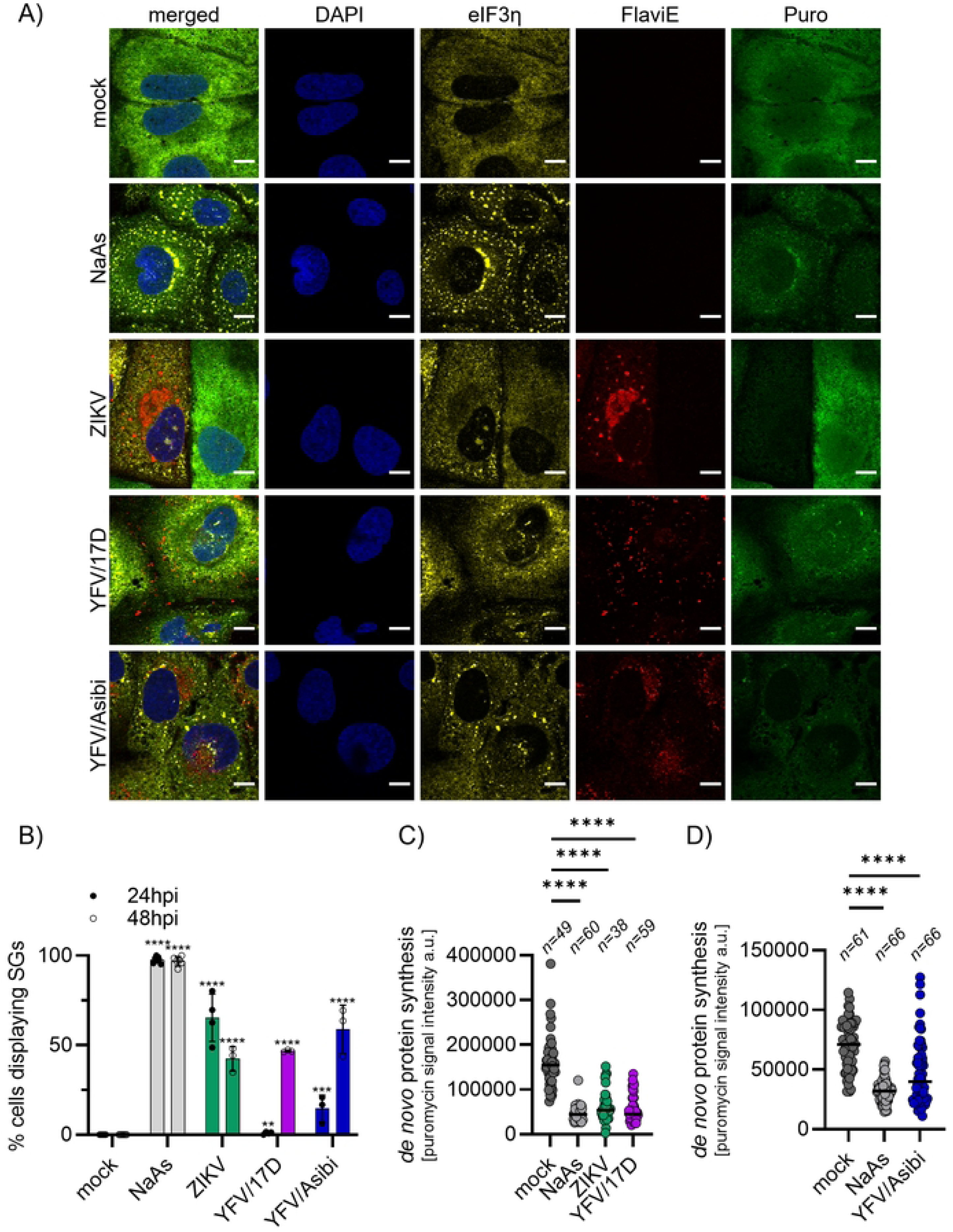
YFV infection results in translational shut-off and SG formation at 48 hpi. Translational activity in virus-infected cells was determined using RPM assays. Naïve Huh7 cells, NaAs-treated cells (50 μM for 30 min) or cells infected with ZIKV, YFV/17D or YFV/Asibi at MOI 1 for 48 h were incubated with 10 μg/ml of puromycin to label nascent peptide chains prior to fixation. Puromycin-labelled peptides were visualized via immunostaining against puromycin (green), infected cells were detected by staining for the flavivirus envelope protein (red) and SGs were observed by staining for eIF3η (yellow). Nuclei were stained with DAPI. **A)** Representative images for the indicated conditions at 48 hpi. Scale bars 10μm. **B)** Quantification of the cell fraction displaying SGs in the conditions indicated at 24 (filled circle) or 48 hpi (open circle). SG formation was assessed by visually inspecting single cells for eIF3η-postive granules. Mock and NaAs-treated cells were included as negative and positive controls respectively. For virus-infected conditions, only cells positive for the flavivirus envelope protein were considered. For each condition at least 170 cells were quantified from at least three biological replicates. Statistical significances relative to mock are indicated at the top. **C)** and **D)** Quantification of translational shut-off for representative replicates for Huh7 cells infected with **C)** ZIKV and YFV/17D or **D)** YFV/Asibi at 48 hpi. Mock and NaAs-treated cells were included as negative and positive controls respectively. Single cells were manually measured for the puromycin signal intensity. For virus-infected conditions, only cells positive for the flavivirus envelope protein were considered. a.u. arbitrary units. Statistical significances and the number of analysed cells (n) are given at the top.

Previous studies have shown that viral infection can induce different SG-like structures, such as RLBs or paracrine granules, which condense SG proteins, such as G3BP1, but form independently of translational activity (21,22). To determine whether the granular structures observed during ZIKV and YFV infection are canonical granules, we treated cells with cycloheximide (CHX). By stabilising polysomes and thereby the shuttling of mRNAs, CHX promotes the disassembly of canonical SGs (35). As a control for canonical SGs we treated cells with silvestrol, an inhibitor of eIF4A helicase activity impairing translation initiation, resulting in polysome disassembly and SG formation (36). Accordingly, while 79 % of silvestrol-stressed cells displayed SGs, upon CHX treatment, only 24 % of cells still contained SGs (Fig 2 A and B). This was also reflected in the decrease in average SG size and number per cell (Fig 2 C and D). CHX treatment of infected cells reduced the number of cells displaying SGs to a comparable extent: for ZIKV from 35 % to 12 % upon CHX addition, for YFV/17D from 42 % to 13 % and for YFV/Asibi from 41 % to 10 % (Fig 2 A and B). To further exclude that the observed granules are RLBs we performed immunostaining for TIA1, which is known to be enriched in canonical SGs, while being excluded from RLBs (S3 Fig) (21). We observed relocalization of TIA1 to ZIKV, YFV/17D and YFV/Asibi-induced granules. Together, this suggests that the granules observed upon infection with YFV are *bona fide* stress granules.

**Fig 2.**
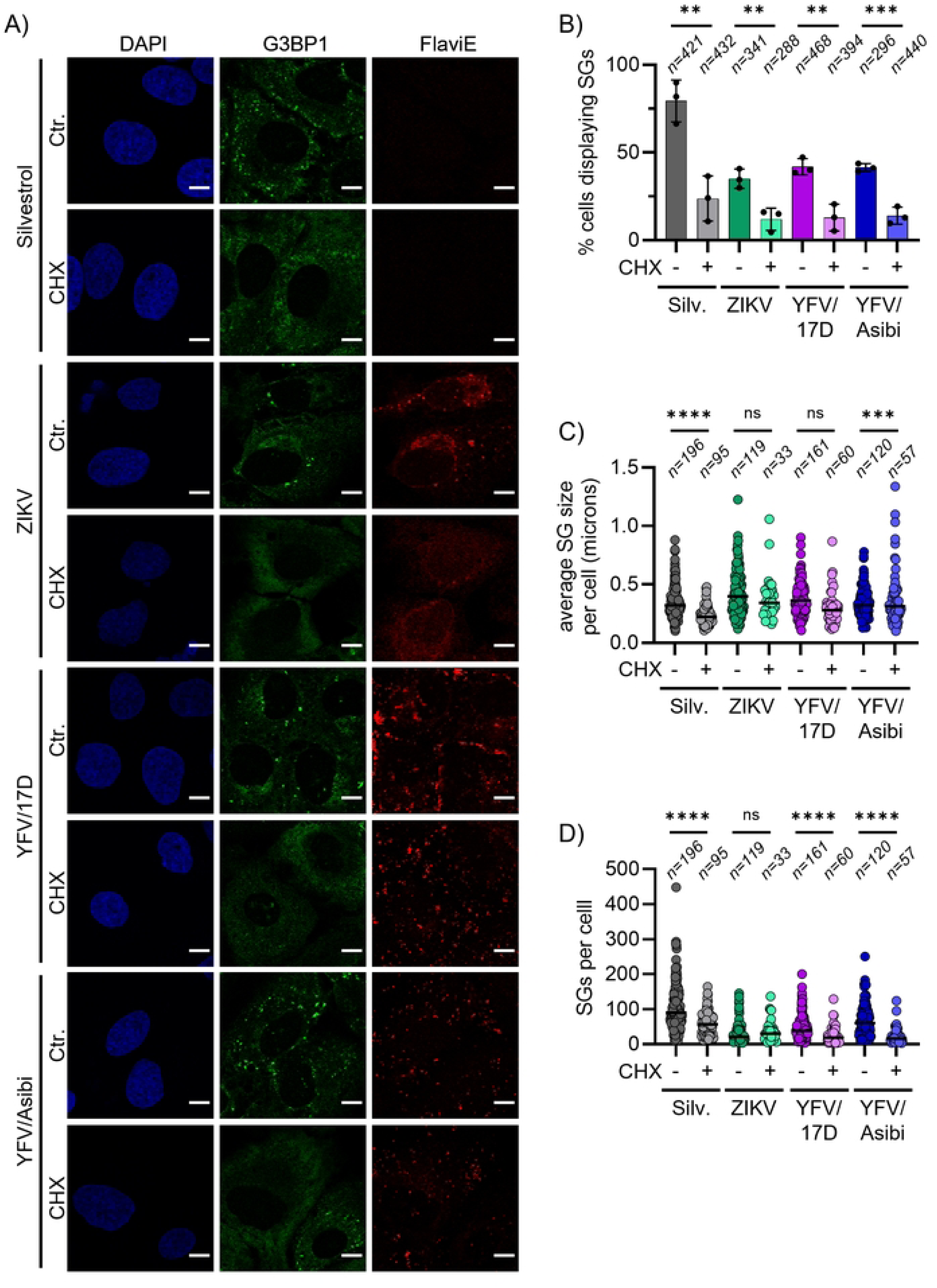
YFV infection induces the assembly of canonical SGs. YFV-induced SGs are sensitive to CHX-mediated disassembly. Huh7 cells infected with ZIKV, YFV/17D or YFV/Asibi at MOI 1 for 48 h were treated with 20 μM CHX for 30 min or left untreated (ctr.) prior to fixation. Silvestrol-treated cells (1 μM for 30 min) were included as positive control for SG formation and CHX-sensitivity. The presence of SGs was visualized via immunostaining against G3BP1 (green), and infected cells were detected by staining for the flavivirus envelope protein (red). Nuclei were stained with DAPI. **A)** Representative images for the indicated conditions at 48 hpi. Scale bars 10μm. **B-D)** SG formation was assessed by visually inspecting single cells for G3BP1-postive granules. For virus-infected conditions, only cells positive for the flavivirus envelope protein were considered. The number of cells analysed (n) from three biological replicates and statistical significances are indicated at the top. **B)** Quantification of the cell fraction displaying SGs. **C)** Measurement of the average SG size in cells displaying SGs. SG size is given in microns. **D)** Quantification of the SG number in cells displaying SGs.

### YFV Capsid interaction might modulate SG nucleation by interaction with G3BP1

A previous study by Hou *et al*. showed that the ectopic expression of the capsid proteins of ZIKV, JEV, MVEV and YFV reduces SG formation in response to chemical stress (27). Since we only observed SG formation in YFV-infected cells at 48 hpi, while translational activity appeared to be repressed already at 24 hpi, we wondered if some of the YFV proteins modulate SG formation. To investigate this, we transfected Huh7 cells with flag-tagged YFV/Asibi proteins and subsequently stressed the cells with NaAs (Fig 3 A and B). Consistent with previous findings, we found that cells transfected with the YFV-Capsid (C) formed SGs less frequently following NaAs treatment compared to untransfected cells. Interestingly, expression of the premature M protein also reduced the number of cells displaying SGs. A previous studies suggested that the ZIKV and JEV capsid protein interacts with the major SG protein G3BP1 (27,32), therefore we tried to identify G3BP1 binding sites within the YFV C protein. Using a motif search algorithm, targeted at identifying binding motifs for the NTF2L-domain of G3BP1 (manuscript in preparation), we predicted a binding site between position 47 and 53 of the YFV-C (Fig 3C, table). To analyse whether this site can interact with G3BP1 we performed fluorescence anisotropy using purified YFV-C peptides and G3BP1-NTF2L domain (Fig 3C, right panel). As a positive control, we used a peptide derived from the SFV-nsp3, since it has previously been demonstrated to interact with G3BP1 (37,38). Similar to the SFV-nsp3 peptide, the YFV-C peptide displayed binding to the G3BP1-NTF2L peptide. Importantly, this binding depended on the predicted G3BP1-NTF2L binding site within the YFV-C, as mutation at position 51 resulted in a diminished interaction. Surprisingly, neither for ZIKV nor the JEV capsid we could predict a NTF2L binding site. Nevertheless, we selected peptides located within the corresponding regions in the ZIKV-C and JEV-C as the identified NTF2L- binding site in YFV-C for fluorescence anisotropy measurement. Neither of the ZIKV-C or JEV-C derived peptides showed any binding to the G3BP1-NTF2L domain. The identification of a G3BP1 binding site within the YFV-C prompted us to check our YFV- SG interactome for the presence of the viral capsid protein. However, none of the YFV proteins were enriched in our proteomics data. Together, these results suggest that individual YFV proteins, such as the YFV-C, can influence SG assembly, potentially via transient interactions with its major nucleator protein G3BP1.

**Fig 3.**
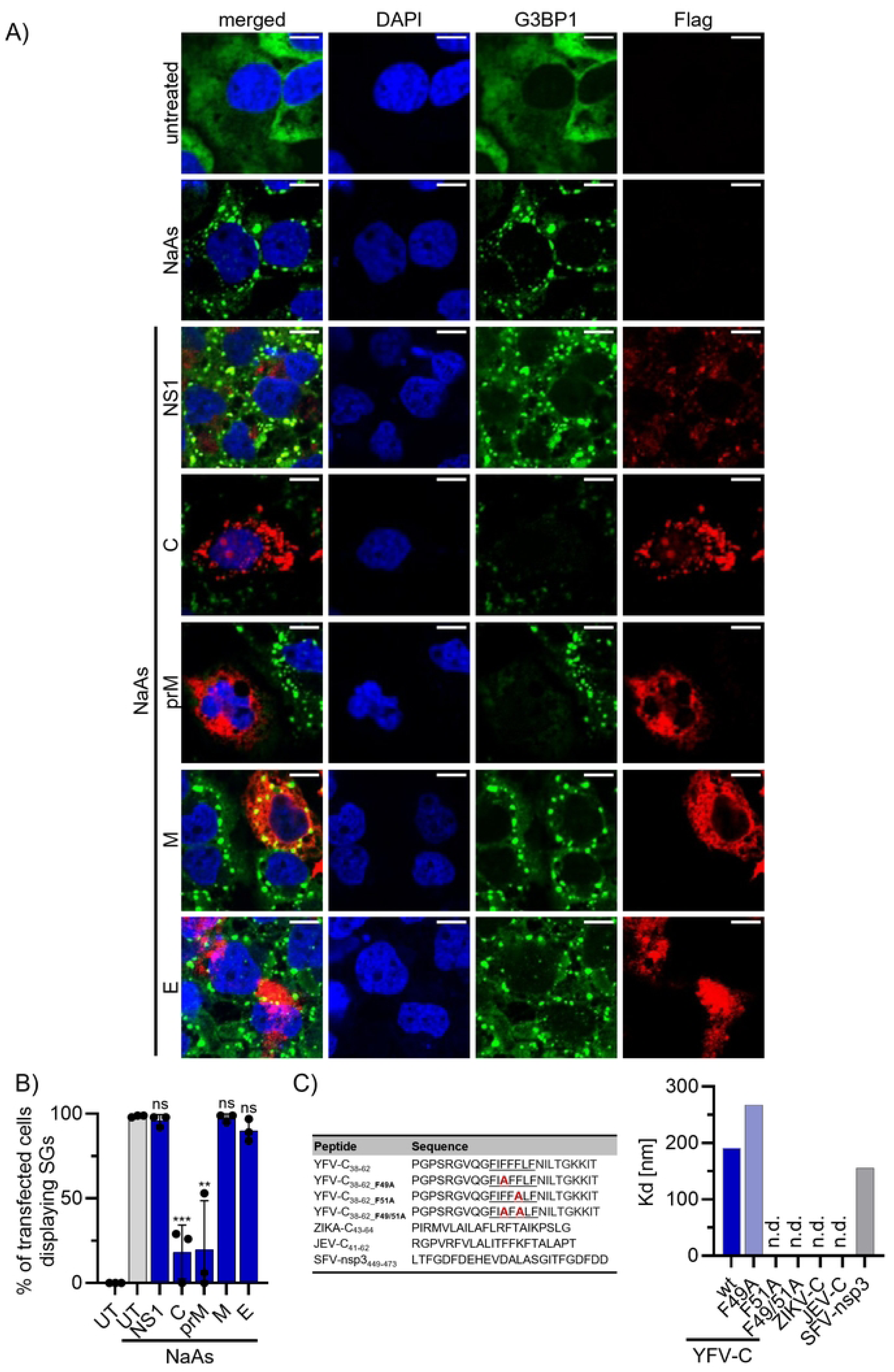
YFV Capsid modulates SG formation and interacts with G3BP1 *in vitro*. **A)** and **B)** Huh7 cells were transfected with plasmids expressing the indicated flag-tagged YFV/Asibi viral proteins or left untransfected (UT) for 48 h. 30 min prior to fixation cells were treated with 50 μM NaAs for 30 min. Transfected cells were identified by immunostaining for the flag tag and the fraction of transfected cells displaying SGs was assessed by staining for G3BP1. **A)** Representative images for the indicated conditions. Scale bars 10μm. **B)** Quantification of Huh7 cells expressing the transfected protein and displaying SGs. Depicted are three biological replicates with the statistical significances given at the top. **C)** Upper panel: Table depicting peptides used to measure NTF2L binding affinity in fluorescence anisotropy binding experiment. YFV-C38-62 contains two overlapping predicted NTF2L-binding motifs (underlined), YFV-C47-51 and YFV-C49-53; F49A, F51A, and F49/51A are mutations that disrupt the binding determinants of the predicted NTF2L-binding motifs (highlighted in bold and red); ZIKA-C43-64 and JEV-C41-62 are located at corresponding regions as the NTF2L-binding site in YFV-C. SFV-nsp3 is a known NTF2L binding peptide. Lower panel: the binding affinity (dissociation constant; Kd) of each peptide in upper panel to the NTF2L domain, measured via fluorescence anisotropy binding experiment.

### Disruption of YFV-induced granules does not impact viral replication

In order to investigate whether YFV-induced granules exhibit anti-viral or pro-viral functions during infection, we sought to disrupt SG formation using a recently identified small-molecule inhibitor, G3Ib (39). G3Ib binds to the main SG scaffolding protein, G3BP1, to block its condensation and thereby disrupts SG formation. The enantiomeric peptide G3Ib’ was used as an inactive control. First, we ensured that treatment with G3Ib efficiently disassembles ZIKV- and YFV-induced granules. For this, cells were infected for 48 h and then treated with G3Ib or the control drug G3Ib’ for 30 min. As a control, cells were treated with silvestrol for 30 min followed by a further 30min treatment with the inhibitor (Fig 4 A, B and C). The active compound G3Ib significantly reduced the number of cells displaying SGs in response to silvestrol from 93 % to 17 % compared to the control drug G3Ib’. Similarly, the fraction of ZIKV- or YFV/Asibi-infected cells forming SGs reduced from 51 % to 24 % or 48 % to 9 % respectively, upon addition of G3Ib. This was also reflected by a reduction in SG number in samples treated with G3Ib (Fig 4 C). Interestingly, G3Ib treatment did not significantly influence the number of cells displaying YFV/17D-induced granules (Fig 4 B), although the it reduced the number of SGs per cell in YFV/17D-infected cells (Fig 4 C). We also assessed the effect of the drugs on cell viability during viral infection, but did not identify any significant differences between the treatments (S4 Fig). Next, we analysed how inhibiting SG formation affects viral replication. To this end, we infected cells with ZIKV, YFV/17D and YFV/Asibi for 48 h in the presence of G3Ib or G3Ib’ and analysed vRNA levels via qPCR and viral titers using plaque assay (Fig 4 D and E). We did not observe a difference in relative vRNA levels or viral titers when SG formation was inhibited in ZIKV- of YFV/Asibi-infected cells. However, for YFV/17D-infected cells, treatment with the active compound G3Ib and the inactive compound G3Ib’ appeared to reduce vRNA levels and viral titers compared to cells treated with the solvent alone. The reasons for this are yet to be defined. These results suggest that the phase-separation of compounds in granules itself does not have a direct pro- or anti-viral effect in the context of YFV infection.

**Fig 4.**
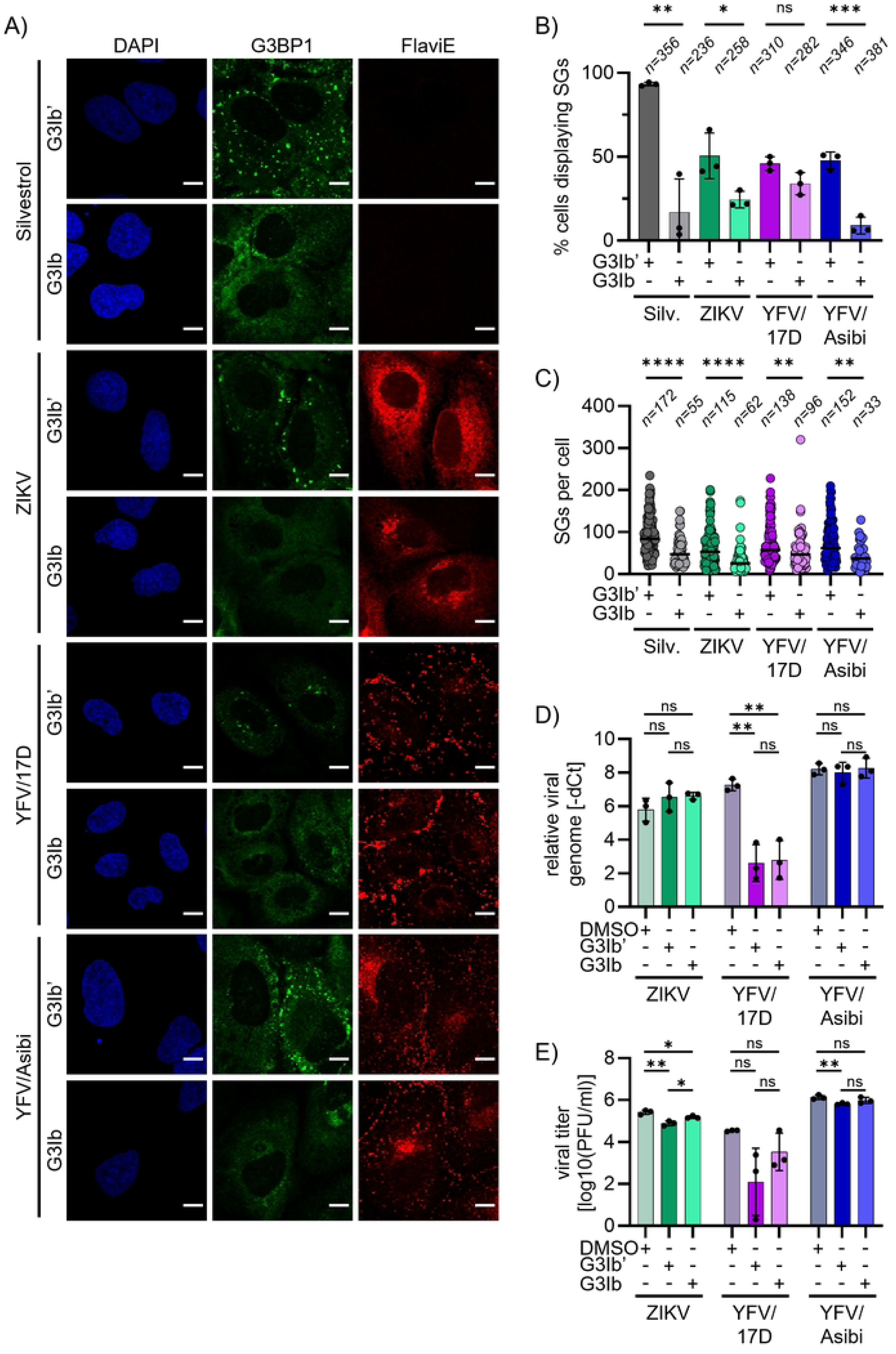
SG disruption does not influence YFV replication. A-C) To prove that the G3BP1-inhibitor G3Ib leads to the disassembly of virus-induced SGs, Huh7 cells infected with ZIKV, YFV/17D or YFV/Asibi at MOI 1 for 48 h were treated with 50 μM G3Ib or the inactive enantiomer G3Ib’ for 30 min prior to fixation. Silvestrol-treated (1 μM for 30 min) Huh7 cells were included as a control for G3Ib-mediated SG disassembly. The presence of SGs was visualized via immunostaining against G3BP1 (green), and infected cells were detected by staining for the flavivirus envelope protein (red). Nuclei were stained with DAPI. **A)** Representative images for the indicated conditions at 48 hpi. Scale bars 10μm. **B-C)** SG formation was assessed by visually inspecting single cells for G3BP1- postive granules. For virus-infected conditions, only cells positive for the flavivirus envelope protein were considered. The number of cells analysed (n) from three biological replicates and statistical significances are indicated at the top. **B)** Quantification of the cell fraction displaying SGs. **C)** Quantification of the SG number in cells displaying SGs. **D-E)** Huh7 cells were infected with ZIKV, YFV/17D or YFV/Asibi at MOI 1 for 48 h in the presence of 50 μM G3Ib or the inactive compound G3Ib’ or DMSO as controls. **D)** Cells were harvested for RNA isolation and subsequent RT-qPCR. Ct values for the viral genome were normalized to the Ct value for the housekeeping gene *tubulin* (-dCT). **E)** Supernatant from infected cells was harvested and viral titer determined via plaque assay. Depicted are the log10-transformed PFU/ml. Statistical significances from three biological replicates are indicated at the top.

### The proteome of YFV-induced stress granules reveals an enrichment of mitochondria-associated proteins

Previous studies have suggested that SGs are heterogenous, adapting stress-specific compositions, which determine their cellular function. Therefore, compositional analysis of SGs have contributed to a better understanding of their function (3,4,21,22,40,41). Accordingly, to elucidate the role of YFV-induced SGs during infection, we aimed to analyse whether these SGs exhibit a different composition compared to NaAs-induced SGs. To do this, we used Huh7 cells that stably express a GFP-tagged G3BP1 and either left these cells untreated (mock) or induced the formation of SGs by the addition of NaAs or infection with YFV/Asibi. Upon cell lysis, the lysate was enriched for SG cores by sequential centrifugation. Next, the granule-containing fraction was purified by immunoprecipitation (IP) using antibodies targeting the GFP-tag of G3BP1 and Protein A-conjugated Dynabeads. The beads were controlled for enrichment of SGs by confirming green fluorescence under the microscope. The resulting protein fraction was analysed by mass spectrometry. We only considered proteins, for which two or more peptides were identified in at least two out of three biological replicates. In all analysed interactomes (mock, NaAs-treated and YFV/Asibi-infected) we identified typical SG proteins such as G3BP2, TIA-1, Caprin1, PABPC1 and UBAP2L (see Sppl. Table 1). While the presence of G3BP1 interactors in untreated cells may seem surprising at first, previous studies have reported that G3BP1 interacts with SG nucleators in unstressed cells, forming ‘seed’ interactions for phase-separation processes upon cellular stress (4). However, in contrast to other studies, in non of the SG proteomes we identified PRRs, such as PKR, RIG-I, MDA-5 or OAS (7–12). As our aim was to identify compositional differences between NaAs- and YFV/Asibi-induced SGs, we selected proteins that were exclusively present in the YFV-SG proteome (Fig 5 A and B). This revealed that 115 proteins are uniquely enriched in YFV-induced SGs. Cellular component (CC) enrichment analysis of these 115 candidates revealed a strong connection to mitochondria, with intrinsic and integral components of mitochondrial membranes being particularly enriched. Similarly, the enriched biological processes included mitochondrial translation and mitochondrial gene expression.

**Fig 5.**
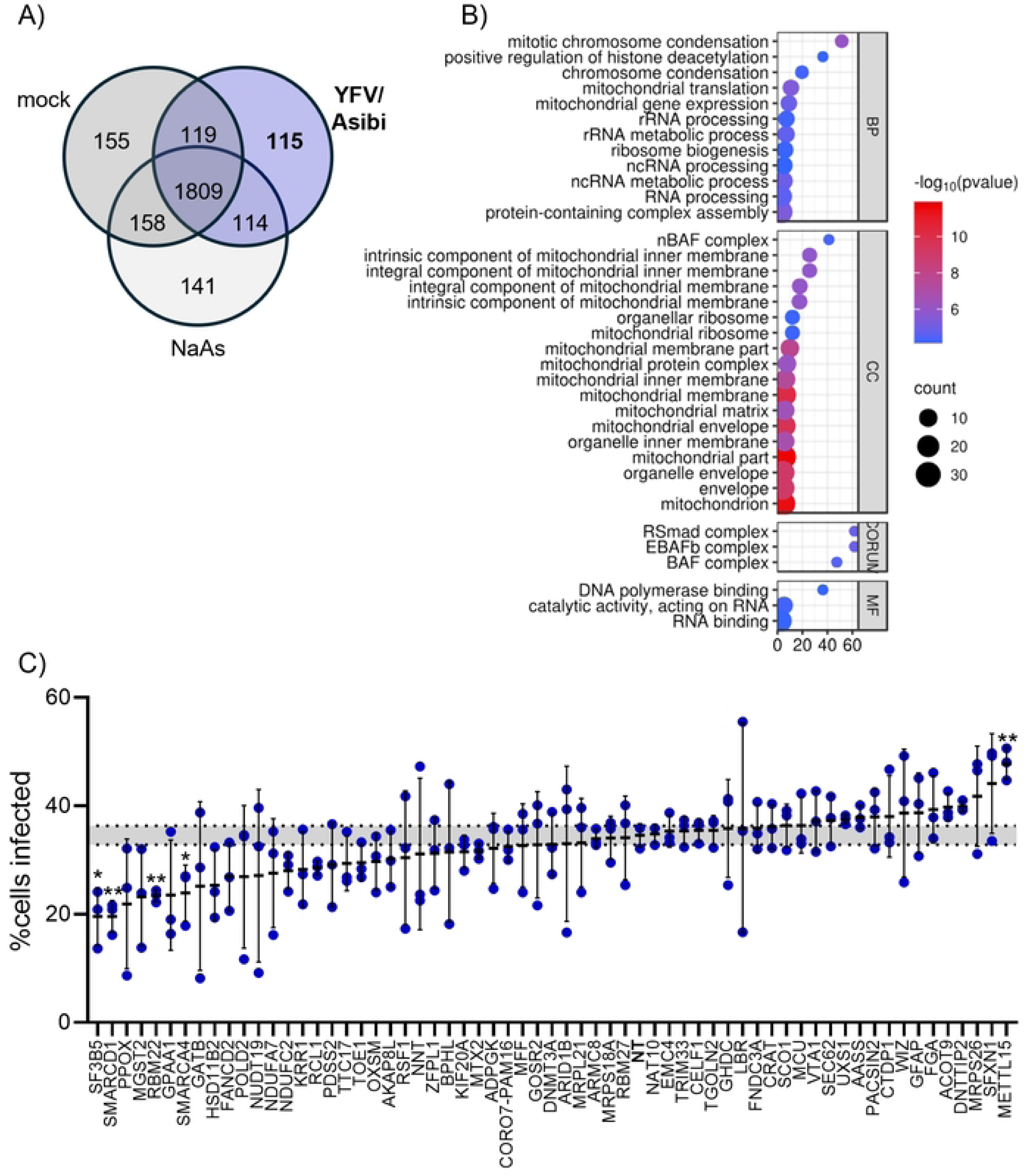
Proteomic analysis of YFV-induced SGs reveals a link to mitochondrial structures and metabolism. **A)** and **B)** GFP-G3BP1 expressing Huh7 cells were either treated with NaAs (0.5 mM for 1 h), infected with YFV/Asibi at MOI 20 or left untreated (mock). At 24 hpi cells were harvested and enriched for the SG-containing fraction, followed by immunoprecipitation for GFP-G3BP1. The resulting eluate was analysed by mass spectrometry. Only proteins for which at least two peptides were found in at least 2 out of 3 replicates were selected. **A)** Venn diagram displaying the number of candidates identified in the indicated conditions**. B)** GO pathway enrichment analysis using WebGestalt (47) of the 115 proteins that were exclusively present in YFV/Asibi-induced SGs. Displayed are most significant terms found in each subset of proteins for biological processes (BP), cellular compartment (CC), comprehensive resource of mammalian protein complexes (CORUM) and molecular function (MF). **C)** siRNA screen of 62 candidates exclusively found in YFV/Asibi-induced SGs. Huh7 cells were transfected with four siRNAs targeting the respective candidate. At 48 h post transfection cells were infected with YFV/Asibi at MOI 20 for 24 h. Infectivity was determined using flow cytometric analysis of cells stained for the viral envelope protein. The percentage of infected cells transfected with a non-target (NT) control is indicated as a grey area. Statistical significances compared to the NT control are indicated at the top.

SG proteins have also been associated with anti- and/or pro-viral functions (reviewed in (18,42)). To investigate whether some of the YFV-SG enriched proteins are involved in modulating viral replication, we selected 62 candidates and performed siRNA screen (Fig 5 C). For this, Huh7 cells were transfected with a set of four siRNAs targeting the respective candidate and subsequently infected with YFV/Asibi. The percentage of infected cells was determined via staining for the flavivirus envelope followed by flowcytometry analysis. While most of the targeted candidates did not influence viral infection rates, knockdown of SF3B5, RBM22, SMARCD1 and SMARCA4 significantly reduced the number of infected cells. SF3B5 and RBM22 are both involved in mRNA splicing, whereas SMARCD1 and SMARCA4 are chromatin remodelers (43–45); therefore, knockdown of these factors might disrupt major cellular processes. Interestingly, knockdown of the mitochondrial methyltransferase METTL15 (46) led to an increase in YFV/Asibi infection, potentially presenting an anti-viral activity of this protein. To summarize, proteomic analysis of YFV/Asibi-induced SGs reveals a distinctive link to mitochondrial structures and potentially function.

### YFV infection leads to mitochondrial damage and dysfunction

As YFV-induced SGs appear to be associated with mitochondrial structures and components, we investigated the impact of YFV infection on mitochondrial shape and function. Previous studies have linked changes in mitochondria morphology to alterations in mitochondrial activity (48). To determine whether YFV infection affects mitochondrial morphology we used a Huh7 cell line that stably expresses a mitochondria-targeted fluorophore (mtTq) (Fig 6 A and B) (49). To control for mitochondrial fragmentation, we treated cells with 50 μM of the uncoupling reagent CCCP for 2 h, which results in mitochondrial damage (50). While 71 % of untreated cells showed tubular mitochondria, CCCP treatment caused mitochondrial fragmentation in the majority of cells. Similarly, infection with YFV/17D or YFV/Asibi reduced the percentage of cells with tubular mitochondria to 7 % and 25 % respectively 48 hpi. In contrast, ZIKV infection appears to influence mitochondrial morphology to a lesser extent, as only 25 % of cells displayed heavily fragmented mitochondria, compared to 74 % for YFV/17D and 42 % for YFV/Asibi-infected cells. Mitochondrial damage is typically associated with an impairment of the electron transport chain resulting in an increase of reactive oxygen species (ROS) and a loss of mitochondrial membrane potential (MMP) (48). To measure ROS levels, we treated cells with MitoSOX reagent, which produces a bright red fluorescence upon oxidation and thereby reflects intracellular ROS levels (Fig 6 C). As a control for ROS production, we treated cells with an uncoupling agent, FCCP at a concentration of 20 μM for 2 h. Compared to mock cells, FCCP-treated as well as ZIKV-, YFV/17D- and YFV/Asibi-infected cells all exhibited increased ROS levels. This correlated with a decrease in MMP in YFV/17D- infected cells from 54 hpi onwards and of YFV/Asibi-infected cells from 57 hpi onwards, as measured using the Incucyte® MMP Orange Reagent (Fig 6 D). As pathway enrichment analysis of YFV-induced SG revealed a link to mitochondrial translation, we aimed to determine the effect of YFV infection on *de novo* mitochondrial protein synthesis. For this, we pulsed the cells with L-homopropargylglycine (HPG) while selectively inhibiting cytosolic translation (51). Thereby, the incorporation rate of HPG into mitochondrial peptides can be used as measurement for mitochondrial translational activity (Fig 6 E and S5 Fig). In cells treated with CCCP, mitochondrial *de novo* protein synthesis was strongly reduced. Similarly, infection with YFV/17D and YFV/Asibi significantly reduced the mitochondrial translational activity. Overall, these data demonstrate that YFV infection results in mitochondrial damage and dysfunction.

**Fig 6.**
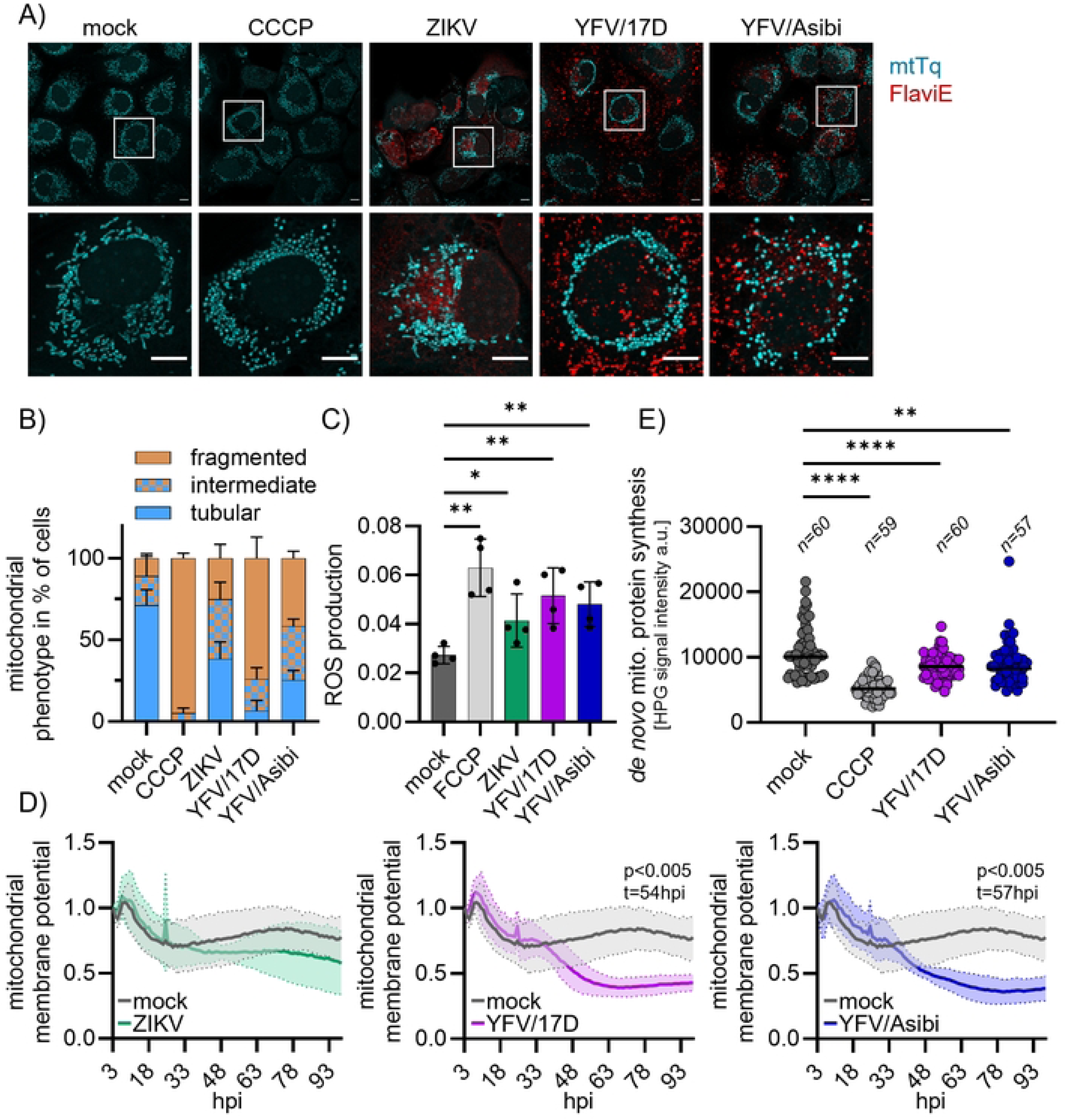
YFV infection leads to mitochondrial damage and dysfunction. **A)** and **B)** Huh7 cells that stably express a mitochondria-targeted fluorophore (mtTq) were infected with ZIKV, YFV/17D or YFV/Asibi at MOI 1 for 48 h. As positive control for mitochondrial fragmentation cells were treated with 50 μM CCCP for 2 hours prior to fixation. Mitochondria are depicted in turquoise; infected cells were detected by immunostaining for the flavivirus envelope protein (red). **A)** Representative images for the indicated conditions at 48 hpi. Scale bars 10μm. Upper panel, overview; lower panel, zoom in. **B)** Quantification of Huh7 cells displaying the indicated mitochondrial phenotypes. For each condition at least 120 cells were quantified from a total of three biological replicates. **C)** ROS levels were determined in naïve Huh7 cells or cells infected with ZIKV, YFV/17D or YFV/Asibi at MOI 2 for 48 h. As control for ROS production cells were treated with 50 μM CCCP for 2 h prior analysis. To measure ROS levels cells were incubated with MitoSoxRed reagent for 30 min prior to determination of the average red object mean intensity by the IncuCyte S3 live cell analysis software. Depicted are four biological replicates with their significances at the top. **D)** Analysis of the mitochondrial membrane potential (MMP) from cells infected with ZIKV, YFV/17D or YFV/Asibi at MOI 2. MMP was monitored for four days every hour using the MMP orange reagent kit and the IncuCyte S3 live cell analysis software. The red mean intensity object average was normalized to 3 hpi. SDs for each time point of five biological replicates are indicated with the shaded area. MMP curves were compared to mock samples using two-way ANOVA with Šídák’s multiple comparisons test. hpi, hours post-infection. **E)** Analysis of *de novo* mitochondrial protein translation. Naïve Huh7 cells, CCCP-treated cells (50 μM for 2 h) or cells infected with YFV/17D or YFV/Asibi at MOI 1 for 48 h were incubated with 50 μg/ml CHX and 1mM HPG to label nascent peptide chains within mitochondria. Upon fixation, HPG was chemoselectively ligated to an azide-containing Alexa Fluor 555 dye. Single cells were manually measured for the HPG signal intensity. Depicted is a representative replicate out of five replicates. a.u. arbitrary units. Statistical significances and the number of analysed cells (n) are given at the top.

## Discussion

While SGs are known to regulate cellular homeostasis and translational control in response to cellular stress, several studies suggest that they take on additional functions during viral infections. It has been proposed that SGs act as a crossroad for the ISR and IIR by concentrating or sequestering PRRs (7–12). However, recent studies challenge this idea, as they have failed to observe the localisation of PRRs to SGs or any adverse effects on IIR activation when the formation of SGs is prevented (10,14–17). Nevertheless, many viruses have evolved different mechanisms to inhibit SG assembly and, or repurpose SG proteins, suggesting that SGs have a function in modulating replication (2,18,19). Molecular dissection of the SG proteome in response to various forms of cellular stress or infection revealed that SGs exhibit stress-specific compositional heterogeneity, which is presumably related to their functional diversity (3,4,21,22,40). In this study, we set out to analyse the cellular stress response to YFV infection and the composition of YFV-induced SGs.

Upon infection with either the YFV/17D vaccine strain or the highly pathogenic YFV/Asibi strain, we observed translational shut-off from 24 hpi onwards, accompanied by canonical SG formation at 48 hpi. Translational shut-off typically coincides with SG formation; therefore, it was surprising to observe that SG induction was delayed by 24 h after translational shut-off became apparent. Interestingly, it has been described for ZIKV and JEV that expression of the viral capsid protein inhibits SG formation (27,32). Similarly, in our study, we found that the ectopic expression of YFV-C in Huh7 cells inhibited NaAs- induced SG formation, as previously reported by Hou *et al* (27). It has been proposed for ZIKV that the viral capsid sequesters the major SG scaffolding protein G3BP1, thereby preventing SG assembly. Using predictive analysis, we identified a G3BP1 binding site at positions 47–53 within YFV capsid, which was confirmed using fluorescence anisotropy. Supporting this finding, a recent preprint by Kenaston *et al*., which analysed the interactome of ectopically expressed and tagged YFV proteins, identified G3BP1 as one of the YFV-C interactors (52). However, we found no viral proteins in the YFV-induced SG interactome, nor did we detect an interaction between G3BP1 and YFV-C in an immunoprecipitation assay against G3BP1 from YFV-infected cell lysates (data not shown). This suggests that the interaction between G3BP1 and YFV-C may be transient and therefore difficult to detect. Nevertheless, during the early stages of infection, the expression of YFV-C may inhibit the formation of SGs. However, as viral replication increases, along with cellular stress, the binding of YFV-C to G3BP1 is insufficient to sequester G3BP1 away from stalled translational complexes, resulting in SG formation. Alternatively, the delayed formation of SGs may correspond to the slow replication kinetics of YFV. A study comparing WNV strains with different infection kinetics suggested that progressing viral replication is associated with a higher tendency to induce SGs (31). Cells infected with a WNV strain producing high vRNA levels early in infection were much more prone to forming SGs than cells infected with a WNV strain exhibiting delayed vRNA synthesis. This was attributed to damage to ER membranes enclosing the viral replication machinery and the subsequent exposure of vRNA to the cytoplasm. Similarly, progressing YFV replication could result in ER membrane damage and vRNA exposure at 48 hpi. This would subsequently be sensed by PKR or RIG-I, as previously reported (17), resulting in eIF2α phosphorylation and SG formation. A combination of the two mechanisms – low vRNA burden during early replication and sequestration of G3BP1 by YFV-C - could potentially contribute to the delayed assembly of SGs during YFV infection.

In this study, we included ZIKV/PRVABC59-infected Huh7 cells as a control for virus induced translational shut-off. While various studies report conflicting results regarding the effect of ZIKV infection on translational shut-off and eIF2α phosphorylation (25–27,30), we clearly observed an increase in eIF2α phosphorylation and reduced translational activity in ZIKV-infected Huh7 cells. However, contrary to previous reports, we observed the assembly of canonical SGs upon ZIKV infection. The differences observed in SG induction and translational shut-off in different experimental set-ups may be related to different viral strains, cell lines, infection loads used, and analysed time points. These factors influence the viral replication burden within infected cells and, consequently, the kinetics of SG formation. This could potentially explain the variable effects of ZIKV infection on the cellular stress response observed in different studies.

To assess the role of SGs during viral infection, previous studies typically employed cell lines depleted of one major SG nucleator protein, such as G3BP1 (18,19,27,53,54). For instance, Bonenfant *et al*. demonstrated that knocking down G3BP1 led to reduced ZIKV replication, whereas knocking down another SG protein, HuR, promoted ZIKV replication (19). In the case of YFV/Asibi, Beauclair *et al*. performed knock-down against four SG nucleator proteins (G3BP1, G3BP2, TIAR and TIA1) to prevent virus-induced SG formation; however, they did not detect any effect on viral replication (17). Although gene silencing provides valuable information on viral dependencies of host cell proteins, it does not address whether the condensation of cellular and/or viral factors into granules influences viral replication *per se*, as it does not allow discrimination between SG-dependent or independent functions of resident proteins. To block SG formation without depleting a particular factor, we used a novel peptide inhibitor called G3Ib, which targets the interaction surface of G3BP1 and thereby prevents SG assembly (39). However, we observed no differences in viral replication or virus-induced cell death in YFV/Asibi- or ZIKV-infected Huh7 cells treated with the drug. This suggests that the formation of the biocondensate itself does not necessarily influence viral replication in the case of YFV/Asibi or ZIKV. However, for YFV/17D, we observed that both G3Ib and its inactive variant negatively affected viral replication, which may suggest adverse effects on yet unidentified cellular mechanisms.

Although we did not observe a role for YFV-induced SG in viral replication, a comparative proteomic analysis of NaAs-induced SG and YFV-induced SG revealed a close link between the viral-induced granules and mitochondrial structures and pathways. siRNA screening of candidates found in YFV-induced granules identified METTL15 as a potential antiviral factor. METTL15 is an N4-methylcytidine (m4C) methyltransferase that is responsible for the methylation of mitochondrial 12S RNA, which is required for mitoribosome biogenesis and mitochondrial translation (46,55). Interestingly, we found that mitochondrial translation was decreased in YFV-infected cells. While the exact manner, in which METTL15 acts as an antiviral factor remains elusive, YFV infection might promote the localisation of METTL15 to SGs, resulting in defective mitochondrial translation.

Interestingly, even components of the inner and intrinsic mitochondrial compartments were enriched in YFV-induced granules. This could be explained in two ways: 1) YFV infection induces mitochondrial damage, and the released components are sequestered in the SG, or 2) YFV-induced SG make close contact with the mitochondria and potentially modulate mitochondrial homeostasis.

Regarding the first hypothesis, we observed that YFV infection induces severe mitochondrial fragmentation, accompanied by increased ROS production, decreased mitochondrial membrane potential, and disrupted mitochondrial translational activity. While preparing this study, a report by Mucilli *et al*. was published describing similar impacts of YFV/17D on mitochondrial activity in HepG2 cells (56). Likewise, previous studies have shown that ZIKV and JEV infection are associated with mitochondrial fragmentation and dysfunction (57–60). For JEV, it has been suggested that mitochondrial fission is necessary for viral replication, as knocking down the mitochondrial fission factor Drp1 reduces viral replication (60). However, conflicting reports exist regarding the effect of DENV on mitochondrial morphology (49,61,62). While Barbier *et al*. and Chatel-Caix *et al*. reported that DENV results in mitochondrial elongation accompanied by dampening of the activation of the IIR, Yu *et al*. observed increased mitochondrial fission in DENV-infected cells. The modulation of mitochondrial activity and fragmentation during ZIKV and JEV infection has been attributed to the expression of the viral NS4a protein, which is targeted to the mitochondria (58,60). However, we could not observe the localisation of the YFV NS4a protein, nor its influence on mitochondrial morphology, when it was ectopically expressed in Huh7 cells (data not shown). Therefore, it is still unclear which YFV protein is responsible for modulating mitochondrial homeostasis.

However, instead of a viral protein, it is possible that the YFV-induced SGs directly influence mitochondrial metabolism via physical contacts. Recent studies have highlighted the functional and physical connections between mitochondria and SGs. Amen *et al*. reported close physical contact between SGs and mitochondria and found that the mitochondrial fatty acid transporter VDAC2 (a transmembrane protein) was enriched in SGs (41). During long-term starvation, SGs appear to downregulate VDAC2, thereby preventing the import of fatty acids into the mitochondria, where they serve as substrates for oxidation and ATP production. It is therefore suggested that SGs reduce fatty acid oxidation in order to save resources and limit ROS production during long-term starvation. We also recently proposed another functional link between the mitochondrial unfolded protein response (mtUPR) and SGs: mtUPR-induced SGs appear to negatively influence mitochondrial homeostasis and cell survival (63). While the interaction between SGs and mitochondrial activity remains largely unexplored, these findings emphasise the close functional relationship between mitochondria and SGs. Similarly, YFV-induced granules may directly affect mitochondrial homeostasis, resulting in fragmentation and dysfunction.

In summary, we described that YFV infection results in activation of the host cell stress response accompanied with translational shut-off and delayed SG formation. While SGs themselves did not appear to influence viral replication, uncovering for the first time the proteome of virus-induced SGs, our analysis revealed a structural and functional link to mitochondria. Indeed, we find that YFV infection results in disruption of mitochondria shape and homeostasis, which is potentially linked to the appearance of SGs upon infection. Overall, this highlights a novel type of virus-host interaction, being the virus-modulated interplay between membrane-bound and membrane-less organelles, contributing to the infection outcome.

## Materials and Methods

### Cell lines and virus strains

Human Huh7 cells (kindly provided by A.Ruggieri, CIID Heidelberg and A. Martin, Institut Pasteur Paris), and ape Vero cells (ECACC, #84113001) were cultured in complete media (Dulbecco’s Modified Eagle Medium (DMEM, Sigma Aldrich #D6429) supplemented with 10 % FBS, 1 % Penicillin-Streptomycin (P/S, Fisher Scientific, #11548876) and 1x L-Glutamine (Fisher Scientific, #11539876)). Huh7 cells stably expressing GFP-tagged G3BP1 (kindly provided by A.Ruggieri, CIID Heidelberg) were cultured in complete media supplemented with sodium pyruvate (Invitrogen). Huh7 mito- mTurquoise2 cells (kindly provided by A.Ruggieri, CIID Heidelberg) were cultured under the addition of 200 ug/ml zeocin (InvivoGen, #ant-zn-1). All cells were grown in a humidified incubator at 37 °C and 5 % CO2. YFV/17D (kindly provided by M. Iqbal, The Pirbright Institute), YFV/Asibi (UVE/YFV/1927/GN/Asibi, EVAg) and ZIKV (PRVABC59, kindly provided by K.Maringer, The Pirbright Institute) virus stocks were grown on Vero cells and the titre determined using plaque assays as described below.

### Virus infection

Huh7 cells were seeded into 24-well or 96-well plates and inoculated the next day with the indicated viruses at the indicated MOI for 2 hours (h) in serum-free DMEM at 37 °C and 5 % CO2. The inoculum was then replaced with complete media (DMEM +10 % FBS +1 % P/S +1 % L-Glutamine). Cells were kept at 37 °C and 5 % CO2 and analysed either at 24 or 48 hours post infection (hpi). For high MOI infections, viruses were concentrated by polyethylene glycol 6000 precipitation and purified by centrifugation in a discontinuous gradient of sucrose.

### Plaque assay

Viral samples were titrated on Vero cells. In short, cells in 12-wells were inoculated with serial dilutions of virus samples in serum-free media at 37 °C and 5 % CO2. After 2h incubation, the inoculum was removed and plaque assay overlay added to the cells. For YFV, either Avicel-containing overlay (DMEM, 1.6 % Avicel (VWR, MANA815290.1), 2 % FBS, 1 % P/S, 1x L-Glutamine) or CMC-containing overlay (1x MEM (Fisher Scientific, #21935-028), 1.5 % CMC (Wako, #039-01335), 1 % FBS, 1 % P/S) was used. For ZIKV, agarose containing overlay was used (DMEM, 1 % LMP Agarose (Invitrogen, #16520- 100), 1 % FBS, 1 % P/S). YFV plaque assays were incubated for 7 days, ZIKV plaque assays for 3 days at 37 °C and 5 % CO2. Plaque assays were fixed using formalin and subsequently stained with toluidine blue.

### qPCR

Huh7 cells were infected in 24 wells with the indicated viruses or left uninfected. At 48 hpi cells were harvested using 200 μl RLT buffer of the RNeasy Mini Kit (Qiagen, #74104) and the RNA isolated according to manufactures protocol. DNA digestion was performed on the column using Turbo DNase (Fisher Scientific, # 10722687) according to manufactures protocol. The RNA was eluted in 30 μl of nuclease-free water and stored at -80 °C. 0.8-1 μg of RNA was reverse transcribed using 200 units of M-MLV Reverse Transcriptase (Promega, #M1705), 0.5 mM dNTPs (Fisher Scientific, # 11853933), 25 units RiboLock RNase Inhibitor (Life Technologies Limited, # EO0382) and 0.5 μg random nanomer primer (Primer Design). For qPCR, the cDNA was diluted 1:100 and 3 μl of the dilution was mixed with 7 μl of PowerUp™ SYBR™ Green Master Mix (Life Technologies Limited, #A25742) and 0.5 μM of forward and reverse primer each (see Sppl. Table 2). qPCR samples were run in technical triplicates in a QuantStudio5 machine. To determine the -dCT value for each target gene, the average Ct value of the technical triplicates was normalised to the respective Ct value of the housekeeping gene *tubulin* as follows: *-dCt=- (Ct_target_-Ct_tubulin_)*

### Ribopuromycylation (RPM) assay

Huh7 cells were seeded on coverslips in 24-wells and infected on the next day with the indicated viruses at MOI of 1 for the indicated times or left uninfected. As positive control for translational shut-off, one well was treated with 0.5 mM sodium arsenite (NaAs) (Fisher Chemical, #S/2330/48) for 30 min. Then, a final concentration of 10 μg/ml puromycin (Merck, P8833) was added to all wells for 5 min at 37°C. Afterwards 180 μM emetine (Merck, #E2375-50MG) was added for 2 min at room temperature prior to fixation with 4 % PFA (Fisher Scientific, #11400580) for 30 min. Immunostaining for eIF3η, puromycin and flavivirus envelope protein was performed as described below. The integrated puromycin signal intensity was determined using the ImageJ software package Fiji by manually fitting ROI for single cells.

### Immunofluorescence assay (IFA)

Fixed cells were permeabilized using 0.1 % Triton-X in PBS for 5 min. After washing with PBS, cells were blocked for 30min with 0.5 % BSA (Sigma, #A3294) in PBS. Samples were subsequently incubated for 1 h with the primary antibody in 0.5 % BSA. After three washes with PBS, the fluorescent labelled secondary antibody together with 0.2 μg/ml DAPI (Invitrogen, #D1306) in 0.5 % BSA was added to the samples for further 1 h. After another three washes with PBS, the coverslips were mounted using 5 μl of Mowiol (Sigma, #91301). Images were acquired using a Leica TCS SP5 Confocal microscope using 40x and 60x objectives.

### Western Blot

Cells in 24 well plates were washed with PBS and harvested in 50 μl 1x Laemmli SDS sample buffer (Fisher Scientific, #15492859c) and heated to 95 °C for 5 min. Samples were loaded onto 4–20 % Mini-PROTEAN® TGX™ Precast Protein Gel (BioRad, #4561096) and subsequently blotted onto nitrocellulose membrane (Cytiva, #10600001) using a Power Blotter Station (Invitrogen). Western blot membranes were blocked with 5 % non-fat dry powder milk in TBS-T (0.2 M Tris-HCl, 1.5 M NaCl, 1 % Tween-20) and subsequently incubated overnight with the respective antibodies (see Sppl. Table 2), either in 5 % milk or, for antibody against phosphorylated antigens, in 5 % BSA. Then membranes were washed with TBS-T and incubated with HRP-coupled anti-mouse or anti-rabbit antibodies (see Sppl. Table 2). Western blot membranes were developed by incubating with Clarity Western ECL substrate (BioRad, #170-5061) and subsequently acquired using a ChemiDoc Imaging System (BioRad). Quantification of western blot bands was performed using the Image Studio Lite Ver 5.2 software.

### Cyclohexamide treatment

Huh7 cells seeded on coverslips in 24-wells were infected as described above with a MOI of 1 or left uninfected. At 48 hpi, a final concentration of 20 μM cyclohexamide (in ethanol, Sigma #C1988-1G) or an equal volume of ethanol (control) was added to infected cells for 30min and incubated at 37 °C. As control, cells were treated with 1 μM silvestrol (MedChemExpress, #HY-13251) for a total of 60 min, while for the last 30 min 20 μM cycloheximide was added to the well. Cells were subsequently fixed with 4 % PFA in PBS for 30 min.

### Cell transfection

Huh7 cells were seeded on coverslips in 24-wells and transfected with pCI-neo-3xFlag plasmids containing the indicated YFV/Asibi-derived sequences using lipofectamine 3000 (Invitrogen, # L3000008) according to manufactures protocol. In short, per transfection, 25 μl mix A (1.5 μl lipofectamine 3000 in OptiMem (Gibco, # 31985062)) was combined with 25 μl mix B (1 μg of the indicated plasmid + 2 μl p3000 reagent in OptiMem) and incubate for 10-15 min at room temperature. Then cell media was exchanged to 250 μl OptiMem and the transfection mixture added dropwise. At 4 h post transfection, media was replaced with complete media. Transfected cells were incubated for 48 h prior to fixation.

### Peptide Synthesis

All peptides were synthesized by the Hartwell Center for Bioinformatics and Biotechnology at St. Jude Children’s Research Hospital, Molecular Synthesis Resource, using standard solid-phase peptide synthesis chemistry (see Sppl. Table 2). All peptides were N-terminal TAMRA-tagged. The peptides were reconstituted from lyophilized form into DMSO for subsequent experiments.

### Recombinant Protein Purification

pGEX-2T-GST-NTF2L construct was transformed into BL21 (DE3) competent cells per the manufacturer’s instructions. Single colony was picked and grown to OD600 of 0.8 in LB medium at 37 °C. Protein expression was then induced with 0.6 mM IPTG overnight at 16 °C. Cells were pelleted by centrifugation at 4,400 x g for 15 min and resuspended in lysis buffer (200 mM NaCl, 50 mM HEPES pH7.5, 1 mM DTT, protease inhibitor). Resuspended cells were lysed by sonication; lysates were pelleted at 30,000 x g for 30 min at 4 °C. Supernatants were applied to GST beads pre-equilibrated with lysis buffer. Proteins were eluted using lysis buffer containing 10 mM glutathione. Eluted proteins were dialyzed in lysis buffer overnight at 4 °C. Dialyzed proteins were collected and concentrated. Protein concentration was determined by OD280 before being flash frozen in liquid nitrogen. Proteins were stored at -80 °C for following experiments.

### Fluorescence Anisotropy Binding Experiments

TAMRA-tagged peptides (100 nM) were incubated with increasing concentrations (50 nM, 100 nM, 200 nM, 400 nM, 800 nM, and 1600 nM) of purified NTF2L in binding buffer (50 mM HEPES pH 7.5, 150 mM NaCl, 1 mM DTT, 0.01 % Tween20) and fluorescence was measured on Horiba/PTI fluorometer with excitation/emission wavelengths of 540/620 nm. GraphPad Prism non-linear fit was used for analysis.

### G3Ib treatment

To check the ability of G3Ib to disassemble virus-induced granules, Huh7 cells seeded on coverslips in 24-wells were infected as described above with a MOI of 1 or left uninfected. At 48 hpi, a final concentration of 50 μM G3Ib or G3Ib’ (39) was added to the cells for 30 min at 37 °C. As control, cells were treated with 1 μM silvestrol for a total of 60 min, while for the last 30 min, 50 μM G3Ib or G3Ib’ was added to the well. Cells were subsequently fixed with 4 % PFA in PBS for 30 min. To assess the effect of granule disassembly on viral replication, Huh7 cells in 24-wells were infected as described above with a MOI of 1 or left uninfected. After virus absorption, complete DMEM containing a final concentration of 50 μM G3Ib or G3Ib’ or an equal volume of DMSO was added to the cells and incubated at 37 °C and 5 % CO2. At 48 hpi, supernatant was harvested for viral titre determination via plaque assay, and cells were harvested for RNA extraction and qPCR as described above.

### Cytotoxicity assay

3x10^3 Huh7 cells were seeded into 96 well plates and inoculated with the indicated viruses at MOI of 2 in serum-free media. After 2 h media was replaced with 200 μl complete media containing 250nM Incucyte® Cytotox Green Dye (Sartorius, # 4633) and the indicated reagents. The plate was sealed using a Breath-Easy sealing membrane (Sigma, #Z380059) and transferred into a Incucyte S3 Live Cell Analysis Instrument. Images of technical triplicates were acquired every 2 h using the 20x objective for a total of 4 days. The green object count per well was determined and normalized to the phase object count per well using the Incucyte 2020A software.

### Stress granule isolation

Stress granules were isolated according to the procedures described in Iadevaia *et al*. (22). Briefly, Huh7 cells stables expressing G3BP1-GFP were grown in 15 cm dishes and subsequently either left uninfected or infected with YFV/Asibi at MOI of 20 by incubation in DMEM containing 2 % FBS for 2h at 37 °C and 5 % CO2. At 24 hpi, cells were harvested and snap frozen until further processing. As control, one plate was treated with 0.5 mM NaAs for 30 min prior to harvesting. The cell pellet was resuspended in 1 ml of lysis buffer (50 mM Tris-HCl, pH 7.4, 100 mM potassium acetate, 2 mM magnesium acetate, 0.5 mM DTT, 50 μg/ml heparin, 0.5 % NP40, cOmplete protease inhibitor (Roche, #11873580001)) using a syringe and a 25G 5/8 needle on ice. Subsequently the cell lysate was centrifuged at 300 g for 5 min at 4 °C. The supernatant was discarded and the pellet again resuspended in 1ml of lysis buffer, followed by another centrifugation step at 18000 g for 20 min at 4 °C. The resulting SG-enriched fraction was again resuspended in 300 μl of lysis buffer and subjected to immunoprecipitation (IP) against GFP. For this, the lysate was precleared using 60 μl of protein A dynabeads (Fisher Scientific #10746713) for 30 min at 4 °C and rotation. Afterwards that lysate was incubated with 0.5 μg anti-GFP antibody (see Sppl. Table 2) at 4 °C and rotation overnight. The lysate was diluted by the addition of 500 μl lysis buffer and subsequently centrifuged again at 18000 g for 20 min at 4 °C. The pellet was resuspended in 500 μl lysis buffer and 33 μl protein A dynabeads and incubated for 3 h at 4 °C and rotation. The beads were washed once with 1 ml of buffer 1 (lysis buffer + 2 M urea) for 2 min, followed by a 5 min wash with 1 ml of buffer 2 (lysis buffer + 300 mM potassium acetate), and another 5 min wash with 1 ml of lysis buffer. Upon seven washed with 1 ml of TE buffer (10 mM Tris, 1 mM EDTA), the beads were sent in TE buffer for MS analysis to the University of Bristol Proteomics Facility.

### Sample preparation for LC-MS/MS analysis

Samples were prepared as described in Iadevaia *et al.* (22). The beads were resuspended in 0.1 M ammonium bicarbonate (ABC) and 0.1 % sodium deoxycholate, then reduced and alkylated using 5 mM TCEP and 20 mM chloroacetamide at 70 °C for 15 min in darkness. Samples were then trypsinized using 0.25 µg of sequencing-grade modified trypsin (Promega) at 42°C for 4 h. Sodium deoxycholate was removed by phase transfer to ethyl acetate. The resulting tryptic peptides were desalted using in-house StageTips with a 3 M Empore SDB-RPS membrane and dried using vacuum centrifugation. The peptides were reconstituted in 15 µl of Buffer A (0.1 % formic acid in water), of which 5 µl was subjected to LC-MS/MS analysis.

### LC-MS/MS analysis

LC-MS/MS analysis was performed as described in Iadevaia *et al*. (22). The tryptic peptides were resolved using a Waters nanoACQUITY UPLC system in a single pump trap mode. The peptides were loaded onto a nanoACQUITY 2G-V/MTrap 5 µm Symmetry C18 column (180 µm×20 mm) with 99.5 % Buffer A and 0.5 % Buffer B (0.1 % formic acid in acetonitrile) at 15 µl/min for 3 min. The trapped peptides were eluted and resolved on a BEH C18 column (130 Å, 1.7 µm×75 µm×250 mm) using gradients of 3 to 5 % B (0–3 min), 8 to 28 % B (3–145 min), and 28 to 40 % B (145–150 min) at 0.3 µl/min. MS/MS was performed on a LTQ Orbitrap Velos mass spectrometer, scanning precursor ions between 400 and 1800 m/z (1×106 ions, 60,000 resolution) and selecting the 10 most intense ions for MS/MS with 180 s dynamic exclusion, 10 ppm exclusion width, repeat count=1, and 30 s repeat duration. Ions with unassigned charge state and MH+1 were excluded from the MS/MS. Maximal ion injection times were 10 ms for Fourier Transform (one microscan) and 100 ms for Linear Ion Trap (LTQ), and the automated gain control was 1×10^4^. The normalized collision energy was 35 % with activation Q 0.25 for 10 ms.

### Mass spectrometry data analysis

Mass spectrometry data analysis was performed as described in Iadevaia *et al*. (22). MaxQuant/Andromeda (version 1.5.2.8) was used to process raw files from LTQ Orbitrap and search the peak lists against the UniProt human proteome database (total 71,803 entries, downloaded 1 December 2018). The search allowed trypsin specificity with a maximum of two missed cleavages and set carbamidomethyl modification on cysteine as a fixed modification and protein N-terminal acetylation and oxidation on methionine as variable modifications. MaxQuant used 4.5 ppm main search tolerance for precursor ions, and 0.5 Da MS/MS match tolerance, searching for the top eight peaks *per* 100 Da. False discovery rates for both protein and peptide were 0.01 with a minimum seven amino acid peptide length.

To define a list of proteins for each experimental condition (mock, NaAs and YFV), we considered the proteins that were identified with at least two peptides, and for which data was obtained in two out of three biological replicates in NaAs and YFV, leaving 2,611 proteins. A total of 155, 115 and 141 proteins were exclusively identified in the mock, YFV and NaAs samples, respectively. Significantly enriched GO terms in YFV (*P* value < 0.01, FDR < 5%) were identified with the GO Term Finder Webgstalt (47). [data will be made publicly available upon acceptance].

### siRNA screen

The ON-TARGETplus siRNA pools used for knockdown (KD) experiments in Huh7 cells were purchased from Horizon (see Sppl. Table 2). Huh7 cells were transfected with Lipofectamine RNAiMAX transfection reagent (Thermo Fisher Scientific, # 13778075) according to the manufacturer’s protocol. 48 h post-transfection, cells were infected with YFV/Asibi at an MOI 10 for 24 h. Subsequently, infected cells were fixed using the Cytofix/Cytoperm fixation and permeabilization kit (BD Pharmingen, # 554722), followed by three washes with the corresponding wash buffer. Subsequently, cells were stained with the flavivirus group antigen antibody (see Sppl. Table 2), diluted in 1 X wash buffer, at 4 °C for 1 h. After another three washes with wash buffer, cells were stained with secondary AF 488 antibody for 45 minutes in the dark at 4 °C. Data acquisition was performed using the Attune NxT Acoustic Focusing Cytometer (Life Technologies), and analysis was conducted using FlowJo v10.8.1 software.

### Mitochondrial fragmentation analysis

Huh7 mito-mTurquoise2 cells seeded on coverslips were infected with the indicated viruses at MOI of 1 for 48 h or left uninfected (mock). As control for mitochondrial fragmentation, cells were treated with 50 μM CCCP (Sigma, #C2759) for 2 h prior to fixation with 4 % PFA in PBS for 30 min. Fixed cells were processed for IFA with the indicated antibodies and confocal pictures acquired as described above. Cells were categorized based on the visual inspection of the mitochondrial shape. If the vast majority of mitochondria appeared as tubular or elongated structures, the cell was categorized as ‘tubular’; if a vast majority of mitochondria appeared as dot-like (fragmented) structures, the cell was categorized as ‘fragmented’; if the same cell contained tubular and fragmented or malformed mitochondria, the cell was categorized as ‘intermediate’.

### Measurement of the mitochondrial reactive oxygen species (ROS) production

3x10^3 Huh7 cells were seeded into 96 well plates and inoculated with the indicated viruses at MOI of 2 or left uninfected (mock) for 48 h. As control, cells were treated with 20 μM FCCP (Sartorius, #6500-0059) for 2 h prior to analysis. For ROS measurement, cell supernatant was removed, cells washed with PBS and 100 μl of PBS containing 2.5 μM MitoSoxRed reagent (Invitrogen, #M36008) added to the wells. After incubation for 30 min at 37 °C, cells were gently washed twice with PBS and 100 μl added to each well. The plate was sealed using a Breath-Easy sealing membrane (Sigma, #Z380059) and transferred into a Incucyte S3 Live Cell Analysis Instrument. Images of technical triplicates were acquired using the 20x objective. The average red object mean intensity, as proxy for ROS levels, was determined using the Incucyte 2020A software.

### Measurement of the mitochondrial membrane potential (MMP)

3x10^3 Huh7 cells were seeded into 96 well plates and inoculated with the indicated viruses at MOI of 2 in serum-free media. After 2 h media was replaced with 200 μl complete media containing 30 nM MMP orange reagent (Sartorius, #4775). The plate was sealed using a Breath-Easy sealing membrane (Sigma, #Z380059) and transferred into a Incucyte S3 Live Cell Analysis Instrument. Images of technical triplicates were acquired every hour using the 20x objective for a total of 4 days. The red mean intensity object average was determined and normalized to 3 hpi using the Incucyte 2020A software.

### Mitochondrial RPM (mtRPM) assay

Huh7 cells were seeded on coverslips in 24-wells and infected on the next day with the indicated viruses at MOI of 1 for 48 h. As positive control for mitochondrial translation shut-off, cells were treated with 50 μM CCCP (Sigma, #C2759) for 2 h. For the mtRPM, media was exchanged with methionine-free media (Sigma, D0422-100ML) and 50 ug/ml cycloheximide (in ethanol, Sigma #C1988-1G), to shut-off cytoplasmic translation, was added to all wells for 20 min at 37 °C. Then, a final concentration of 1 mM Click-iT® homopropargylglycine (HPG) (Invitrogen, # C10186) was added to the wells and incubated for 20 min at 37 °C. Then, the plate was put on ice and cells washed with PBS, followed by the addition of cold 500 μl of pre-permeabilization buffer (10 mM Hepes, 10 mM NaCl, 5 mM KCl, 300 mM sucrose) containing 0.015 % digitonin for 2 min. Afterwards, cells were washed with pre-permeabilization buffer for 15 seconds and subsequently fixed with 4 % PFA in PBS for 30 min. The click-reaction addition of AZDye555-Picolyl-Azide (Jena Bioscience, # CLK-1288-AZ) to HPG was performed using the CuAAC Cell Reaction Buffer Kit (BTTAA based) (Jena Bioscience, # CLK-073) according to manufactures protocol. In short, samples were washed with PBS and incubated with 100 mM NH4Cl for 15 min, followed by incubation with 5 % BSA in PBS for 10 min. A final concentration of 9.1 mM of CuSO4 and 45.45 mM BTTAA were pre-mixed together. Per sample a click-reaction solution containg 168.75 μl of 100 mM Na-Phosphate reaction buffer, pH 7, 20 μl of 500 μM AZDye555-Picolyl-Azide, 55 μl of the CuSO4:BTTAA premix and 25 μl of 1 M Na-Ascorbate stock solution was mixed together and subsequently added to the coverslip. Samples were incubated for 60 min in the dark, followed by two washes with 5 % BSA in PBS. Then, samples were processed for IFA against the mitochondrial marker TOMM20 (see Sppl. Table 2) as described above. The integrated HPG signal intensity was determined using the ImageJ software by manually fitting ROI for single cells.

### Analysis of SG frequency, average SG size and SG numbers per cell

Confocal pictures were analysed using the ImageJ software package Fiji v.1.50. To determine the frequency of cells displaying SGs, cells were manually counter using the multi-point plugin. For infected conditions, only cells that showed staining for the flavivirus envelope protein were considered. To determine the average SG size and SG number per cell, first ROIs were fitted manually for single cells. Then, the threshold on the channel for the SG marker was manually adjusted to mask the punctuated SG signal. Using the Image J ParticleAnalyzer plugin (https://imagej.net/ij/developer/api/ij/ij/plugin/filter/ParticleAnalyzer.html) the average SG size and the SG number was determined for each ROI.

### Statistical analysis

Statistical analysis and data visualisation was done using GraphPad Prism version 10.2.0 for Windows, GraphPad Software, Boston, Massachusetts USA, www.graphpad.com. If not indicated otherwise, data was analysed using unpaired t test. Significances are indicated as follows: ns=p>0.05; *=p≤0.05; **= p≤0.01; ***= p≤0.001; ****= p≤0.0001.

## Acknowledgement

We sincerely thank to Alessia Ruggieri (CIID, Heidelberg, Germany), Kevin Maringer, Jean-Remy Sadeyen, Munir Iqbal (The Pirbright Institute, Woking, UK) for providing cell lines and viruses. Further we would like to thank the Pirbright Institute CL3 core facility (BBSRC NBRI grant: BBS/E/PI/23NB0004) for providing BSL3 training and the Bioimaging core facility (BBSRC NBRI grant: BBS/E/PI/23NB0003) for technical assistance with the confocal microscopy.

## References

1. Hofmann S, Kedersha N, Anderson P, Ivanov P. Molecular mechanisms of stress granule assembly and disassembly. Biochim Biophys Acta Mol Cell Res. 2021 Jan;1868(1):118876.

2. Brownsword MJ, Locker N. A little less aggregation a little more replication: Viral manipulation of stress granules. Wiley Interdiscip Rev RNA. 2023;14(1):e1741.

3. Jain S, Wheeler JR, Walters RW, Agrawal A, Barsic A, Parker R. ATPase modulated stress granules contain a diverse proteome and substructure. Cell. 2016 Jan 28;164(3):487–98.

4. Markmiller S, Soltanieh S, Server KL, Mak R, Jin W, Fang MY, et al. Context-Dependent and Disease-Specific Diversity in Protein Interactions within Stress Granules. Cell. 2018 Jan 25;172(3):590–604.e13.

5. Niewidok B, Igaev M, Pereira da Graca A, Strassner A, Lenzen C, Richter CP, et al. Single-molecule imaging reveals dynamic biphasic partition of RNA-binding proteins in stress granules. J Cell Biol. 2018 Apr 2;217(4):1303–18.

6. Streicher F, Jouvenet N. Stimulation of Innate Immunity by Host and Viral RNAs. Trends Immunol. 2019 Dec;40(12):1134–48.

7. Reineke LC, Lloyd RE. The stress granule protein G3BP1 recruits protein kinase R to promote multiple innate immune antiviral responses. J Virol. 2015 Mar;89(5):2575–89.

8. Onomoto K, Jogi M, Yoo JS, Narita R, Morimoto S, Takemura A, et al. Critical role of an antiviral stress granule containing RIG-I and PKR in viral detection and innate immunity. PLoS One. 2012;7(8):e43031.

9. Yoo JS, Takahasi K, Ng CS, Ouda R, Onomoto K, Yoneyama M, et al. DHX36 enhances RIG-I signaling by facilitating PKR-mediated antiviral stress granule formation. PLoS Pathog. 2014 Mar;10(3):e1004012.

10. Langereis MA, Feng Q, van Kuppeveld FJ. MDA5 localizes to stress granules, but this localization is not required for the induction of type I interferon. J Virol. 2013 Jun;87(11):6314–25.

11. Manivannan P, Siddiqui MA, Malathi K. RNase L Amplifies Interferon Signaling by Inducing Protein Kinase R-Mediated Antiviral Stress Granules. J Virol. 2020 Jun 16;94(13):e00205–20.

12. Paget M, Cadena C, Ahmad S, Wang HT, Jordan TX, Kim E, et al. Stress granules are shock absorbers that prevent excessive innate immune responses to dsRNA. Mol Cell. 2023 Apr 6;83(7):1180–1196.e8.

13. Fujikawa D, Nakamura T, Yoshioka D, Li Z, Moriizumi H, Taguchi M, et al. Stress granule formation inhibits stress-induced apoptosis by selectively sequestering executioner caspases. Curr Biol. 2023 May 22;33(10):1967–1981.e8.

14. Zappa F, Muniozguren NL, Wilson MZ, Costello MS, Ponce-Rojas JC, Acosta- Alvear D. Signaling by the integrated stress response kinase PKR is fine-tuned by dynamic clustering. J Cell Biol. 2022 May 6;221(7):e202111100.

15. Burke JM, Ratnayake OC, Watkins JM, Perera R, Parker R. G3BP1-dependent condensation of translationally inactive viral RNAs antagonizes infection. Sci Adv. 10(5):eadk8152.

16. Corbet GA, Burke JM, Bublitz GR, Tay JW, Parker R. dsRNA-induced condensation of antiviral proteins modulates PKR activity. Proc Natl Acad Sci U S A. 2022 Aug 16;119(33):e2204235119.

17. Beauclair G, Streicher F, Chazal M, Bruni D, Lesage S, Gracias S, et al. Retinoic Acid Inducible Gene I and Protein Kinase R, but Not Stress Granules, Mediate the Proinflammatory Response to Yellow Fever Virus. J Virol. 2020 Oct 27;94(22):e00403–20.

18. Jayabalan AK, Griffin DE, Leung AKL. Pro-Viral and Anti-Viral Roles of the RNA- Binding Protein G3BP1. Viruses. 2023 Feb 6;15(2):449.

19. Bonenfant G, Williams N, Netzband R, Schwarz MC, Evans MJ, Pager CT. Zika Virus Subverts Stress Granules To Promote and Restrict Viral Gene Expression. J Virol. 2019 Jun 15;93(12):e00520–19.

20. Loucas G, Locker N, Parker R. Nucleic acid-protein condensates in innate immunity. Molecular Cell. 2025 Oct 16;85(20):3823–39.

21. Burke JM, Lester ET, Tauber D, Parker R. RNase L promotes the formation of unique ribonucleoprotein granules distinct from stress granules. J Biol Chem. 2020 Feb 7;295(6):1426–38.

22. Iadevaia V, Burke JM, Eke L, Moller-Levet C, Parker R, Locker N. Novel stress granule-like structures are induced via a paracrine mechanism during viral infection. J Cell Sci. 2022 Feb 15;135(4):jcs259194.

23. Genus: Orthoflavivirus | ICTV [Internet]. [cited 2025 Sep 25]. Available from: https://ictv.global/report/chapter/flaviviridae/flaviviridae/orthoflavivirus

24. Douam F, Ploss A. Yellow fever virus: Knowledge gaps impeding the fight against an old foe. Trends Microbiol. 2018 Nov;26(11):913–28.

25. Roth H, Magg V, Uch F, Mutz P, Klein P, Haneke K, et al. Flavivirus Infection Uncouples Translation Suppression from Cellular Stress Responses. mBio. 2017 Jan 10;8(1):e02150–16.

26. Ricciardi-Jorge T, da Rocha EL, Gonzalez-Kozlova E, Rodrigues-Luiz GF, Ferguson BJ, Sweeney T, et al. PKR-mediated stress response enhances dengue and Zika virus replication. mBio. 2023 Oct 31;14(5):e0093423.

27. Hou S, Kumar A, Xu Z, Airo AM, Stryapunina I, Wong CP, et al. Zika Virus Hijacks Stress Granule Proteins and Modulates the Host Stress Response. J Virol. 2017 Aug 15;91(16):e00474–17.

28. Courtney SC, Scherbik SV, Stockman BM, Brinton MA. West Nile Virus Infections Suppress Early Viral RNA Synthesis and Avoid Inducing the Cell Stress Granule Response. Journal of Virology. 2012 Apr;86(7):3647–57.

29. Tu YC, Yu CY, Liang JJ, Lin E, Liao CL, Lin YL. Blocking Double-Stranded RNA- Activated Protein Kinase PKR by Japanese Encephalitis Virus Nonstructural Protein 2A. Journal of Virology. 2012 Oct;86(19):10347–58.

30. Amorim R, Temzi A, Griffin BD, Mouland AJ. Zika virus inhibits eIF2α-dependent stress granule assembly. PLoS Negl Trop Dis. 2017 Jul;11(7):e0005775.

31. Emara MM, Brinton MA. Interaction of TIA-1/TIAR with West Nile and dengue virus products in infected cells interferes with stress granule formation and processing body assembly. Proc Natl Acad Sci U S A. 2007 May 22;104(21):9041–6.

32. Katoh H, Okamoto T, Fukuhara T, Kambara H, Morita E, Mori Y, et al. Japanese Encephalitis Virus Core Protein Inhibits Stress Granule Formation through an Interaction with Caprin-1 and Facilitates Viral Propagation. J Virol. 2013 Jan;87(1):489–502.

33. Hahn CS, Dalrymple JM, Strauss JH, Rice CM. Comparison of the virulent Asibi strain of yellow fever virus with the 17D vaccine strain derived from it. Proc Natl Acad Sci U S A. 1987 Apr;84(7):2019–23.

34. Taniuchi S, Miyake M, Tsugawa K, Oyadomari M, Oyadomari S. Integrated stress response of vertebrates is regulated by four eIF2α kinases. Sci Rep. 2016 Sep 16;6:32886.

35. Kedersha N, Cho MR, Li W, Yacono PW, Chen S, Gilks N, et al. Dynamic Shuttling of Tia-1 Accompanies the Recruitment of mRNA to Mammalian Stress Granules. J Cell Biol. 2000 Dec 11;151(6):1257–68.

36. Cencic R, Carrier M, Galicia-Vázquez G, Bordeleau ME, Sukarieh R, Bourdeau A, et al. Antitumor Activity and Mechanism of Action of the Cyclopenta[b]benzofuran, Silvestrol. PLOS ONE. 2009 Apr 29;4(4):e5223.

37. Panas MD, Ahola T, McInerney GM. The C-Terminal Repeat Domains of nsP3 from the Old World Alphaviruses Bind Directly to G3BP. Journal of Virology. 2014 May 15;88(10):5888–93.

38. Panas MD, Varjak M, Lulla A, Eng KE, Merits A, Karlsson Hedestam GB, et al. Sequestration of G3BP coupled with efficient translation inhibits stress granules in Semliki Forest virus infection. Mol Biol Cell. 2012 Dec;23(24):4701–12.

39. Freibaum BD, Messing J, Nakamura H, Yurtsever U, Wu J, Kim HJ, et al. Identification of small molecule inhibitors of G3BP-driven stress granule formation. J Cell Biol. 2024 Mar 4;223(3):e202308083.

40. Brocard M, Iadevaia V, Klein P, Hall B, Lewis G, Lu J, et al. Norovirus infection results in eIF2α independent host translation shut-off and remodels the G3BP1 interactome evading stress granule formation. PLoS Pathog. 2020 Jan;16(1):e1008250.

41. Amen T, Kaganovich D. Stress granules inhibit fatty acid oxidation by modulating mitochondrial permeability. Cell Rep. 2021 Jun 15;35(11):109237.

42. Guan Y, Wang Y, Fu X, Bai G, Li X, Mao J, et al. Multiple functions of stress granules in viral infection at a glance. Front Microbiol. 2023 Mar 1;14:1138864.

43. Sun C. The SF3b complex: splicing and beyond. Cell Mol Life Sci. 2020 Mar 5;77(18):3583–95.

44. Soubise B, Jiang Y, Douet-Guilbert N, Troadec MB. RBM22, a Key Player of Pre- mRNA Splicing and Gene Expression Regulation, Is Altered in Cancer. Cancers (Basel). 2022 Jan 27;14(3):643.

45. Centore RC, Sandoval GJ, Soares LMM, Kadoch C, Chan HM. Mammalian SWI/SNF Chromatin Remodeling Complexes: Emerging Mechanisms and Therapeutic Strategies. Trends in Genetics. 2020 Dec 1;36(12):936–50.

46. Van Haute L, Hendrick AG, D’Souza AR, Powell CA, Rebelo-Guiomar P, Harbour ME, et al. METTL15 introduces N4-methylcytidine into human mitochondrial 12S rRNA and is required for mitoribosome biogenesis. Nucleic Acids Res. 2019 Nov 4;47(19):10267–81.

47. Elizarraras JM, Liao Y, Shi Z, Zhu Q, Pico AR, Zhang B. WebGestalt 2024: faster gene set analysis and new support for metabolomics and multi-omics. Nucleic Acids Res. 2024 Jul 5;52(W1):W415–21.

48. Cervantes-Silva MP, Cox SL, Curtis AM. Alterations in mitochondrial morphology as a key driver of immunity and host defence. EMBO reports. 2021 Sep 6;22(9):e53086.

49. Chatel-Chaix L, Cortese M, Romero-Brey I, Bender S, Neufeldt CJ, Fischl W, et al. Dengue Virus Perturbs Mitochondrial Morphodynamics to Dampen Innate Immune Responses. Cell Host & Microbe. 2016 Sep 14;20(3):342–56.

50. Miyazono Y, Hirashima S, Ishihara N, Kusukawa J, Nakamura K ichiro, Ohta K. Uncoupled mitochondria quickly shorten along their long axis to form indented spheroids, instead of rings, in a fission-independent manner. Sci Rep. 2018 Jan 10;8(1):350.

51. Yousefi R, Fornasiero EF, Cyganek L, Montoya J, Jakobs S, Rizzoli SO, et al. Monitoring mitochondrial translation in living cells. EMBO reports. 2021 Apr 7;22(4):e51635.

52. Kenaston MW, Cherkashchenko L, Skawinski CLS, Fishburn AT, Peddamallu V, Florio CJ, et al. Yellow Fever Virus Interactomes Reveal Common and Divergent Strategies of Replication and Evolution for Mosquito-borne Flaviviruses [Internet]. bioRxiv; 2025 [cited 2025 Oct 8]. p. 2025.06.14.659623. Available from: https://www.biorxiv.org/content/10.1101/2025.06.14.659623v2

53. Scholte FEM, Tas A, Albulescu IC, Žusinaite E, Merits A, Snijder EJ, et al. Stress granule components G3BP1 and G3BP2 play a proviral role early in Chikungunya virus replication. J Virol. 2015 Apr;89(8):4457–69.

54. Zheng ZQ, Wang SY, Xu ZS, Fu YZ, Wang YY. SARS-CoV-2 nucleocapsid protein impairs stress granule formation to promote viral replication. Cell Discov. 2021 May 25;7(1):1–11.

55. Chen H, Shi Z, Guo J, Chang KJ, Chen Q, Yao CH, et al. The human mitochondrial 12S rRNA m4C methyltransferase METTL15 is required for mitochondrial function. J Biol Chem. 2020 Jun 19;295(25):8505–13.

56. Muccilli SG, Schwarz B, Jessop F, Shannon JG, Bohrnsen E, Shue B, et al. Mitochondrial Hyperactivity and Reactive Oxygen Species Drive Innate Immunity to the Yellow Fever Virus-17D Live-Attenuated Vaccine. bioRxiv. 2024 Sep 15;2024.09.04.611167.

57. García CC, Vázquez CA, Giovannoni F, Russo CA, Cordo SM, Alaimo A, et al. Cellular Organelles Reorganization During Zika Virus Infection of Human Cells. Front Microbiol. 2020;11:1558.

58. Lee JK, Shin OS. Zika virus modulates mitochondrial dynamics, mitophagy, and mitochondria-derived vesicles to facilitate viral replication in trophoblast cells. Front Immunol. 2023;14:1203645.

59. Yang S, Gorshkov K, Lee EM, Xu M, Cheng YS, Sun N, et al. Zika Virus-Induced Neuronal Apoptosis via Increased Mitochondrial Fragmentation. Front Microbiol. 2020;11:598203.

60. Agarwal A, Alam MF, Basu B, Pattanayak S, Asthana S, Syed GH, et al. Japanese Encephalitis Virus NS4A Protein Interacts with PTEN-Induced Kinase 1 (PINK1) and Promotes Mitophagy in Infected Cells. Microbiol Spectr. 2022 Jun 29;10(3):e0083022.

61. Barbier V, Lang D, Valois S, Rothman AL, Medin CL. Dengue virus induces mitochondrial elongation through impairment of Drp1-triggered mitochondrial fission. Virology. 2017 Jan;500:149–60.

62. Yu CY, Liang JJ, Li JK, Lee YL, Chang BL, Su CI, et al. Dengue Virus Impairs Mitochondrial Fusion by Cleaving Mitofusins. PLoS Pathog. 2015 Dec;11(12):e1005350.

63. Lopez-Nieto M, Sun Z, Relton E, Safakli R, Freibaum BD, Taylor JP, et al. Activation of the mitochondrial unfolded protein response regulates the dynamic formation of stress granules. J Cell Sci. 2025 May 1;138(9):jcs263548.

